# A mechanochemical model recapitulates distinct vertebrate gastrulation modes

**DOI:** 10.1101/2021.10.03.462928

**Authors:** Mattia Serra, Guillermo Serrano Nájera, Manli Chuai, Vamsi Spandan, Cornelis J. Weijer, L. Mahadevan

## Abstract

Gastrulation is a critical event in vertebrate morphogenesis driven by cellular processes, and characterized by coordinated multi-cellular movements that form the robust morphological structures. How these structures emerge in a developing organism and vary across vertebrates remains unclear. Inspired by experiments on the chick, we derive a theoretical framework that couples actomyosin activity to tissue flow, and provides a basis for the dynamics of gastrulation morphologies. Our model predicts the onset and development of observed experimental patterns of wild-type and perturbations of chick gastrulation as a spontaneous instability of a uniform state. Varying the initial conditions and a parameter in our model, allows us to recapitulate the phase space of gastrulation morphologies seen across vertebrates, consistent with experimental observations in the accompanying paper. All together, this suggests that early embryonic self-organization follows from a minimal predictive theory of active mechano-sensitive flows.

---

Gastrulation is an essential, highly conserved process in the development of all vertebrate embryos (1), with the chick being an extensively studied model because it is easily cultured and imaged. During gastrulation, the chick transforms from a layer of epithelial cells into a layered structure of three major embryonic tissues, the ectoderm, mesoderm and endoderm. At the moment of egg-laying, the chick embryo contains around 60,000 cells organized in a central concentric epiblast that will give rise to the embryo proper (EP) surrounded by a ring of extraembryonic (EE) tissue. During the first few hours of development, signals from the extraembryonic tissues, the epiblast and the developing hypoblast induce mesendoderm precursor cells located in a sickle-shaped domain at the posterior edge of the epiblast (2) (Fig. 1A). These mesendoderm precursor cells start to execute directed cell intercalations that results in a contraction of the mesendoderm tissue towards the central midline followed by an extension in anterior direction, forming the primitive streak (PS) (3,4) (Figs. 1A-B). In the streak, the mesendoderm precursor cells undergo an epithelial to mesenchymal transition; they ingress individually and migrate away to form the various meso and endodermal structures and organs (5, 6). This directional cell intercalation process drives embryo scale counter-rotating tissue flows in the epiblast merging at the site of the formation of the PS (7, 8) (Fig. 1B).

**Figure 1:**
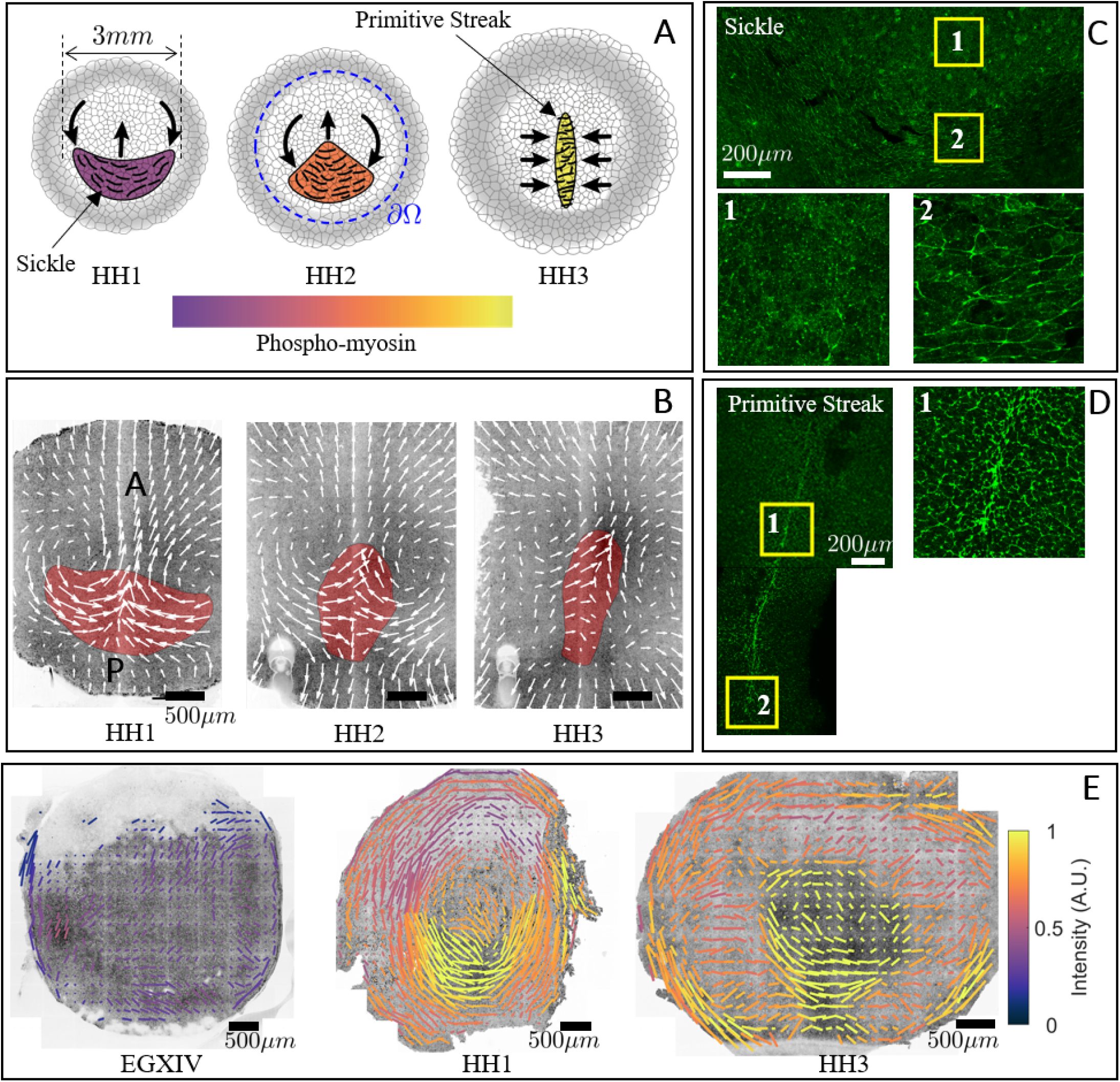
Active myosin cables drive the tissue flows in gastrulation. A) Diagram of the formation of the PS. B) Formation of the PS on a chick gastrula. The convergent extension of the mesendoderm (red) originate macroscopic tissue flows on the surface of the embryo. C) The mesendoderm sickle territory is characterised by supracellular cables of active myosin (phospho-myosin light chain, pMLC) perpendicular to the anterior-posterior (AP) direction. The PS is characterised by supracellular cables of active myosin (pMLC) perpendicular to the midline. E) The pattern of actomyosin cables evolves over time. The bars represent the pMLC anisotropy and the color of the bars indicates the relative concentration. For details see the accompanying paper (9).

There are three cellular scale active force-generating processes that drive tissue flows: (i) the outward migration of the cells attached to the vitelline membrane at the boundary of the embryo, (ii) the active intercalation of the mesendoderm precursors generating forces in the plane of the epiblast, (iii) the ingression of mesendoderm cells in the streak, wich attracts cells towards the streak and drives the out of plane motion. It is likely that the variation of tissue flows and morphologies seen during gastrulation across vertebrates, ranging from fish and amphibians via reptiles to amniotes such as chick and human, are due to changes in the embryo geometry and the relative contribution of these three force-generating processes.

At the cellular level, in the chick, the onset of directional intercalation correlates with the appearance of oriented chains of aligned junctions containing active actomyosin, as detected by phosphorylation of the myosin light chain (4, 10) (Figs. 1A left, C). This indicates the appearance of a supra-cellular oriented organization (planar cell polarity) of intercalating cells (SFig. 4). During the extension of the primitive streak the supracellular cables of highly active myosin reorient to become perpendicular to the midline (Figs. 1A right, D, E). These myosin cables, first observed in directional cell intercalation underlying germband extension in fruitfly embryogenesis (11, 12), were thought to arise from signaling in the anterior-posterior patterning system (13). More recent work (14) suggests instead that the myosin cables could self-organize in a tension-dependent manner, and thus lead to spontaneous large-scale orientation during PS formation. Consistent with this, it is now well established that cytoskeletal actin dynamics, myosin activity and adhesion, are mechano-sensitive (15), e.g. contraction of a given junction will increase its tension, which in turn activates the myosin assembly and further myosin recruitment through a catch-bound mechanism (16–18) (Fig. 2A). This positive feedback mechanism could then result in the formation of supra-cellular chains of myosin-enriched junctions at the cellular level that drive coordinated supra-cellular directional intercalation, organized tissue flow and PS formation (2). In the chick, the early embryonic layer can be abstracted as a thin two-dimensional fluid in which myosin activity leads to active stresses which induce multicellular flows. Interfering with myosin activity correlates with the failure of PS formation (4) and accompanying paper (9). But what are the essential mechanisms sufficient to generate supra-cellular coordination?

**Figure 2:**
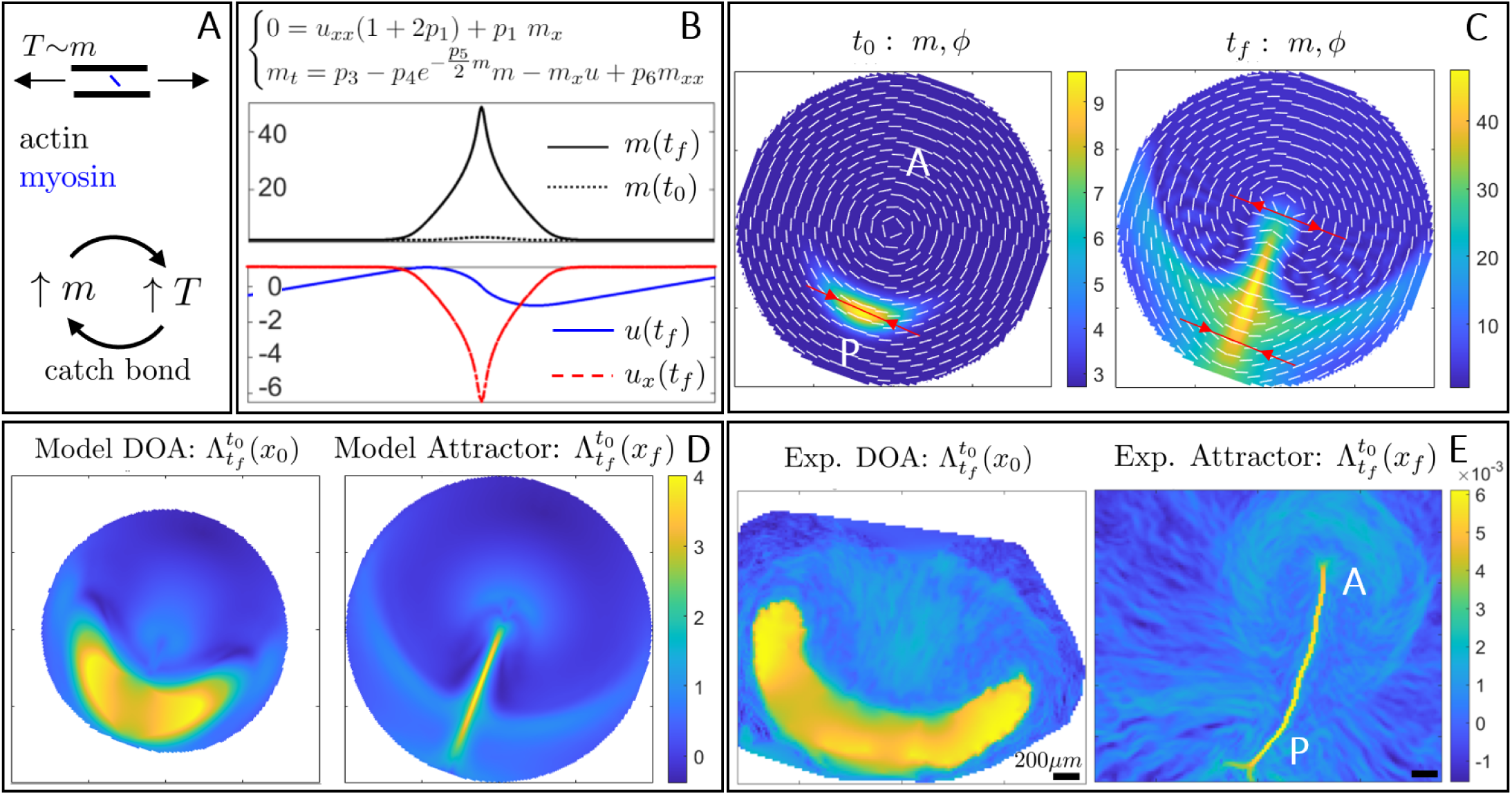
Dynamics of wild-type gastrulation in the chick. A) High tension *T* along actomyosin cables induces additional myosin recruitment via the catch-dynamics, further increasing *T*. This positive feedback process causes the instability of eq. (1). B) 1D Model recapitulates a focusing-type instability of active myosin, which induces a co-located velocity sink describing cell ingression at the PS. [Movie1](https://www.dropbox.com/s/6pgxys02xe2or9e/1D%20Model.mp4?dl=0) shows the time evolution of *m, u, u*_*x*_. C) Initial (*t*_0_) and final (*t*_*f*_) distributions of *m, ϕ* for the 2D model. The instability of *m* drives the redistribution of cables orientation generating compression (tension) at the P (A) PS. D) Model-based DOA at *t*_0_ and the attractor at *t*_*f*_. [Movie2](https://www.dropbox.com/s/n3s5ylkqqd4nv2s/WT.mp4?dl=0) shows the time evolution of the relevant model-based Eulerian and Lagrangian fields. E) Same as D from the experimental velocity. [Movie3](https://www.dropbox.com/s/z4xhncptk04wfk3/DynMor_CTR0245_M1Exp.mp4?dl=0) shows the time evolution of the Lagrangian metrics from the experimental **v**. The colorbar in E shows attraction rates in min^−1^. See materials and methods for values of parameters and boundary conditions used in simulating Eq. (1).

To probe the conditions for the emergence and self-organization of ordered active myosin cables during development and correlate these with observed tissue flows in normal and perturbed conditions, we need a quantitative predictive model. Minimally, a planar mechanochemical model of PS formation during gastrulation should couple three coarse-grained fields: the tissue velocity field **v**(*x, y, t*) = (*u*(*x, y, t*), *v*(*x, y, t*))^⊤^, the active stress intensity *m*(*x, y, t*) arising from the active myosin, and the cable orientation *ϕ*(*x, y, t*). In the limit of slow viscously-dominated flows associated with morphogenesis, we can neglect inertia so that local force balance reads ∇ *· σ*_*T*_ = **0**, where the total stress *σ*_*T*_ = *σ*_*V*_ + *σ*_*A*_ is the sum of the viscous stress *σ*_*V*_ = *−p***I** + 2*μ***S**_*s*_ (*μ* being the shear viscosity and **S**_*s*_ = (∇**v** + ∇**v**^*T*^ *−* (∇ *·* **v**)**I**)*/*2 the deviatoric rate-of-strain tensor), and the active stress, 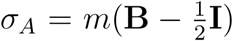, where **B** = **e** ⊗ **e** characterizes the local stress tensor associated with the contractile myosin cables whose orientation vector is given by **e** = (cos *ϕ*, sin *ϕ*)^⊤^.

Our model of the sheet-like embryo is assumed to be a compressible fluid to accommodate cell ingression. This implies that we need a continuity law that accounts for flow divergence (or convergence). A simple linear model for this is ∇ *·* **v** = *c* (Trace[*σ*_*T*_] *− p*_0_*m*). Here *c*^*−*1^ is the fluid bulk viscosity while *p*_0_ characterizes the effect of *active cell ingression* and can be thought of as the ratio between the isotropic to anisotropic active stress (see material and methods for a detailed description and the implications of this law), and reflects the expectation that both the average compressive stress and active myosin contribute to negative divergence and thence cell ingression. Then, we may write the equations for local momentum balance for the planar velocity field **v**(*x, y, t*), along with equations for the orientational *ϕ*(*x, y, t*) and intensity *m*(*x, y, t*) dynamics of myosin cables as

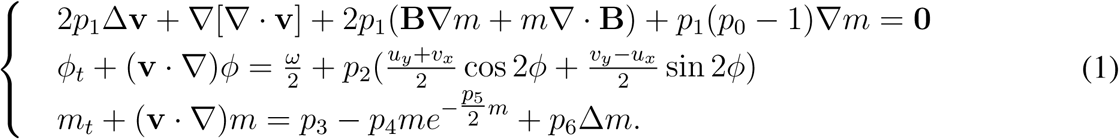

Here we have nondimensionalized our model using the characteristic length scale *x*_*c*_, speed *u*_*c*_ and viscous shear stress *μu*_*c*_*/x*_*c*_. In the first equation in (1), the first two terms are the contributions from shear and dilatational deformations, the last three terms are a consequence of active forces induced by inhomogeneities in the myosin concentration and cable orientation, while *p*_1_ = *μc* is the ratio of the shear viscosity to the bulk viscosity (see materials and methods § 1-2 for a derivation of these equations). The second equation in (1) describes the orientational dynamics of myosin cables which evolve like material fibers, advected by the flow and rotated by vorticity *ω* and shear (19), with the non-dimensional parameter *p*_2_ *∼ O*(1) for elongated fibers. The last equation tracks the magnitude of the active stress intensity associated with the recruitment of myosin from the cytoplasm, assumed to have a uniform constant concentration *m*_*n*_, at scaled rate 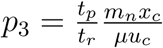 (here *t*_*p*_ is the phosphorylation time scale for converting active myosin into active stress and *t*_*r*_ is the recruitment time scale). The second term on the right of the last equation is associated with the positive feedback induced by the catch-bond dynamics in loaded cables (Fig. 2A) that has an exponential form (20) (with *m/*2 being the cable tension and 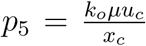 is the ratio of the characteristic shear stress of the system to the characteristic bonding stress between actin and myosin). 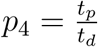 is the ratio of phosphorylation time scale to the dissociation time scale, and the last term accounts for myosin diffusion with 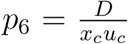 being the inverse of the Peclet number characterizing the ratio between diffusive and advective transport of myosin. Given the circular geometry of the embryo, we solve our model in polar coordinates (see materials and methods for a detailed derivation, nondimensionalization, discussion and numerical scheme). We consider a spatial domain with boundary *∂*Ω (Fig. 1A) that encloses the embryonic area and a small fraction of the EE region. This allows us to keep the size of the domain approximately fixed, as the embryonic area remains constant. To complete the formulation of the problem, we need boundary and initial conditions. For boundary conditions, we impose the epiboly velocity and no flux for *m, ϕ*, i.e. **v** = **v**_b_|_*∂*Ω_, ∇*m ·* **n** = **0**, ∇*ϕ ·* **n** = **0**, with **n** being the outer normal to *∂*Ω. See materials and methods for the selection of the parameters in our simulations.

To gain intuition, we derive and analyze (materials and methods) the 1D version of eq. (1) modeling the dynamics perpendicular to AP, as summarized in Fig. 2B. Linear stability analysis of uniform equilibria of *m* reveals that the (lower) higher equilibrium is linearly (stable) unstable (SFig. 2). Initializing the nonlinear system with a Gaussian perturbation *m*(*t*_0_) to the unstable equilibrium (mimicking the onset of actomyosin cables in the sickle as in Figs.1A,C), our model develops a focusing-type instability which significantly increases *m*(*t*_*f*_) while generating a highly compressive region *u*_*x*_(*t*_*f*_) ≪ 0, reproducing cell ingression at the streak. [Movie1](https://www.dropbox.com/s/6pgxys02xe2or9e/1D%20Model.mp4?dl=0) shows the time evolution of the relevant fields. While the simplified 1D model neglects the orientation of the cables, it accounts for the catch-bond dynamics coupled with the flow field and our continuity law, both of which carry over the full model.

To quantify the global nature of the self-organized spatio-temporal patterns, we use the notion of Lagrangian coherent structures adapted to morphogenetic flows, i.e. their dynamic morphoskeletons (DM) (21). The DM is based on a Lagrangian description of tissue deformation captured by the Finite Time Lyapunov Exponents (FTLE), which naturally combines local and global mechanisms along cell paths. The DM consists of attractors and repellers toward which cells converge or diverge over a specific time interval *T* = *t*_*f*_ *− t*_0_. Repellers are marked by high values of the forward FTLE 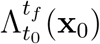; attractors are marked by high values of backward FTLE 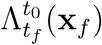 and their domain of attraction (DOA) by high values of the backward FTLE displayed on the initial cell positions 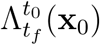 (see materials and methods for details).

We initialize the 2D model with a curved Gaussian perturbation (mimicking the mesendoderm precursor) to the unstable equilibrium of *m* and *ϕ*(*t*_0_) in the azimuthal direction (Fig. 2C), consistent with experiments (Fig. 1E and Fig. 3 of (9)). The instability of *m* drives both the flow velocity and the orientation of the cables shown in Fig. 2C and [Movie2](https://www.dropbox.com/s/n3s5ylkqqd4nv2s/WT.mp4?dl=0), reproducing the typical flow patterns observed in wild-type experiments. A closer look at *m, ϕ* shows that the active stress resulting from our self-organizing model induces tension in the anterior and compression in the posterior (red arrows Fig. 2C), resembling the stress distribution on the tensile ring observed in (10). While (10) imposes the tensile ring on a fixed circle located at the boundary between the EP and EE area, our active stress distribution arises spontaneously from Eq. (1) and evolves dynamically in space. This is consistent with our experimental results where we identify the EP-EE boundary as a repeller (21) and track its spatio-temporal evolution (SFig. 5 and [SM Movie1](https://www.dropbox.com/s/yslitzz0wwjon92/OuterRepEvolution_CTR0245_M2Exp.mp4?dl=0)). Figures 2D-E show the DOA (left) and the attractor (right), corresponding to the largest *T* obtained from both the model and the experimental **v**. The attractor marks the formed PS, while the DOA marks the initial position of cells that will form the PS (see [Movie2](https://www.dropbox.com/s/n3s5ylkqqd4nv2s/WT.mp4?dl=0) for the time evolution of the model-based *m, ϕ*, **v**, attractors, repellers and a deforming Lagrangian grid as *T* increases, and [Movie3](https://www.dropbox.com/s/z4xhncptk04wfk3/DynMor_CTR0245_M1Exp.mp4?dl=0) for the same Lagrangian quantities obtained from the experimental velocity fields). These results complement Fig. 1A-G and [Movie1](https://www.dropbox.com/s/6pgxys02xe2or9e/1D%20Model.mp4?dl=0) of (9) showing brightfield images and gene expression patterns that indicate the cable orientation and myosin concentration, consistent with our model. All together, a comparison of the model- and experiment-based Lagrangian metrics shows that our model accurately explains the appearance of the localized zone of myosin activity and the concomitant flows as a spontaneous instability that leads to a self-organized dynamical structure, in contrast to its imposition at a fixed location between the EP and EE in earlier work (10).

**Figure 3:**
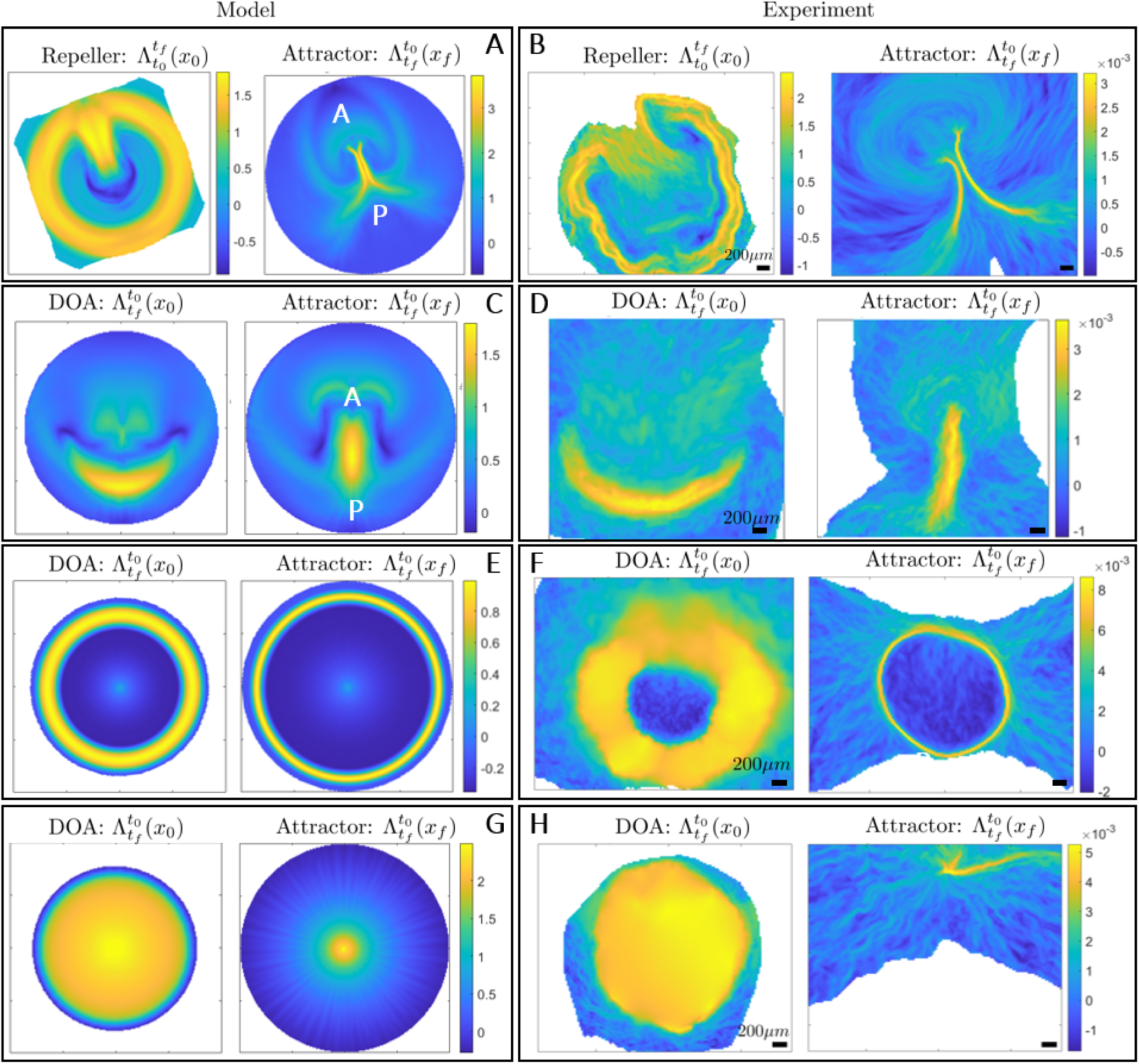
Developmental perturbations of gastrulation. A-B) Spontaneous twin perturbation. A) Model-based repeller and attractor for the largest *T*. [Movie4](https://www.dropbox.com/s/mesc5gvojmtofr1/PTB_TWIN.mp4?dl=0) shows the time evolution of the relevant model-based Eulerian and Lagrangian metrics. B) Same as A from the experimental **v**. [Movie5](https://www.dropbox.com/s/k4q4dtywyd5frlz/InAndFinTime_FWBWFTLE__PTB0475.mp4?dl=0) shows the time evolution of the Lagrangian metrics from the experimental **v**. C-D) Inhibition of cell ingression at the PS. C) Model-based DOA and attractor for the largest *T*. [Movie6](https://www.dropbox.com/s/s8b8lajdf97in2q/PTB_FGF_Rot.mp4?dl=0) shows the time evolution of the relevant model-based Eulerian and Lagrangian metrics. D) Same as C from the experimental **v**. [Movie7](https://www.dropbox.com/s/1va949bys6514w5/InAndFinTime_FWBWFTLE__PTB0449_Rot.mp4?dl=0) shows the time evolution of the Lagrangian metrics from the experimental **v**. E-F) FGF2 addition provokes a circular PS. E) shows the same as C. [Movie8](https://www.dropbox.com/s/qidwjcbo75zjxl2/PTB_CRC.MP4?dl=0) shows the same as Movie7. F) and [Movie9](https://www.dropbox.com/s/e10su20anexviu8/InAndFinTime_FWBWFTLE__PTB0552_v2.mp4?dl=0) are the same as E and Movie8 from the experimental **v**. G-H) BMP and GSK3 induce a ring-shaped mesoderm territory at the EE-EP interface while blocking apical contraction and cell ingression in the PS. G) shows the same as C. [Movie11](https://www.dropbox.com/s/vj56wmwsglw170k/PTB_CRC_FGF.mp4?dl=0) shows the same as Movie7. H) and [Movie12](https://www.dropbox.com/s/k7&796mocto4jmr/InAndFinTime_FWBWFTLE__PTB0505.mp4?dl=0) are the same as G and Movie11 from the experimental **v**. Colorbars in experimental panels show attraction rates in min^−1^. See materials and methods for values of parameters and boundary conditions used in simulating Eq. (1).

To further test the predictive power of our model, we move beyond gastrulation in wild-type avian embryos, and introduce four different perturbations that modify either the initial state of the embryonic pattern (e.g. its mesoderm) or the active cell ingression encoded in *p*_0_. Occasionally, a natural event corresponding to the spontaneous formation of twins arises when the mesendoderm precursor area is split in two regions, from which two streaks emerge. These streaks interact through their tissue flows and form complete or partially twined embryos (Figs. 3A-B). We initialize our model as in the wild type (Fig. 2C) but adding two distant Gaussians to the uniform unstable equilibrium of *m*. [Movie4](https://www.dropbox.com/s/mesc5gvojmtofr1/PTB_TWIN.mp4?dl=0) shows the model-based Eulerian fields and the induced Lagrangian metrics for increasing *T*. Figure 3A shows the model-based repeller and the attractor for the largest *T*. The circular repeller separates the EP from the EE region as well as the AP regions of the PS, while the attractor marks the merging PSs. Figure 3B and [Movie5](https://www.dropbox.com/s/k4q4dtywyd5frlz/InAndFinTime_FWBWFTLE__PTB0475.mp4?dl=0) show the same as 3A and Movie 4 for the experimental **v**. Comparison of Movie 4 and 5 highlights how our model recapitulates the full morphogenetic process quantified clearly by the DM.

In the second perturbation, we interfere with a signalling pathway via application of the VEGF receptor (Vascular Epithelial Growth Factor) inhibitor Axitinib (100nM), which results in an strong inhibition of ingression of cells in the streak (Figs. 3C-D). We initialize our model as in the wild type (Fig. 2C) but set *p*_0_ = 0 as Axitinib prevents active cell ingression. Figures 3C-D show the DOA and attractor for both the model and experimental **v**, highlighting how this perturbation results in a shorter and thicker PS as well as a reduced amount of ingressed cells, consistent with the results shown in Fig. 2H-N and [Movie6](https://www.dropbox.com/s/s8b8lajdf97in2q/PTB_FGF_Rot.mp4?dl=0) of (9). [Movie6](https://www.dropbox.com/s/s8b8lajdf97in2q/PTB_FGF_Rot.mp4?dl=0) and [Movie7](https://www.dropbox.com/s/1va949bys6514w5/InAndFinTime_FWBWFTLE__PTB0449_Rot.mp4?dl=0) show the time evolution of the model- and experiment-based fields confirming a strikingly similar DM again. The third perturbation consists of FGF2 addition which provokes the generation of a mesoderm ring along the marginal zone that essentially behaves like a circular PS. For this case, we use the same conditions of the wild type, but set *m*(*t*_0_) as a circular ring added to the unstable equilibrium (see *m*(*t*_0_) in [Movie8](https://www.dropbox.com/s/qidwjcbo75zjxl2/PTB_CRC.MP4?dl=0)). Figures 3E-F show the DOA and attractor corresponding to the largest *T* from the model and experimental **v**, highlighting a sharp circular PS. [Movie8](https://www.dropbox.com/s/qidwjcbo75zjxl2/PTB_CRC.MP4?dl=0) and [Movie9](https://www.dropbox.com/s/e10su20anexviu8/InAndFinTime_FWBWFTLE__PTB0552_v2.mp4?dl=0) show the Eulerian fields and the Lagrangian metrics for increasing *T*, while [Movie10](https://www.dropbox.com/s/5mawtinn5nd8fpz/fgf2_expk0552_grid.avi?dl=0) shows a deforming Lagrangian grid overlaid on the light-sheet microscope images. These results complement those in Fig. 1H-N of (9) and confirm the predictive power of our model. As the last perturbation, we induce the formation of mesoderm with a combination of the BMP (Bone Morphogenetic Protein) receptor inhibitor LDN*−*193189 (100nM) and the GSK3 (Glycogen Synthase Kinase 3) inhibitor CHIR*−*99021 (3*μ*M). This induces a ring shaped mesoderm territory at the EE-EP interface. This treatment furthermore blocks apical contraction and cell ingression in the PS, but has little effect on cell intercalations, resulting in the buckling of the tissue (Fig. 2A-G and [Movie 5](https://www.dropbox.com/s/mi91ky38e11l8w8/movie_05_LDN_CHIR.avi?dl=0) of (9)). In our model, we use the same setting of perturbation 3 but set *p*_0_ = 0 following the same reasoning of perturbation 2. Figures 3G-H show the DOA and the attractor for to the largest *T* from the model and the experimental **v**, while [Movie11](https://www.dropbox.com/s/vj56wmwsglw170k/PTB_CRC_FGF.mp4?dl=0) and [Movie12](https://www.dropbox.com/s/k7&796mocto4jmr/InAndFinTime_FWBWFTLE__PTB0505.mp4?dl=0) display the corresponding time evolution of the Eulerian fields and Lagrangian metrics.

To go beyond studying the gastrulation modes in wild-type avian embryos exemplified by the chick, in the accompanying paper (9) we used chemical inhibitors to uncouple cell intercalation and ingression and perturb the initial patterning and function of the prospective mesendoderm. Our results show that we can recapitulate some of the major evolutionary transitions in the morphodynamics of vertebrate gastrulation. The current paper complements this in terms of a mechanochemical model that couples the dynamics of actomyosin cables to large-scale tissue flow patterns. By changing the isotropic to anisotropic active stress ratio, characterized by the parameter *p*_0_, together with the initial prospective mesendoderm, described by *m*(*t*_0_), (Fig. 4 left), we can capture the observed variations in the modes of vertebrate gastrulation (Fig. 4 center, right). Specifically, the PS structures in Figs. 3 C-D resemble the blastoporal groove observed in reptilian gastrulation, the circular streak structure (Figs. 3 E-F) resembles the germband of teleost fish gastrulation, while the buckling tissue in Figs. 3 G-H mimics the tissue organization and flow during the closing blastopore in amphibians. Altogether, these results suggest that a relatively small number of changes in critical cell behaviors associated with different force-generating processes leads to distinct vertebrate gastrulation modes via a self-organizing process that might be relatively easily evolvable.

**Figure 4:**
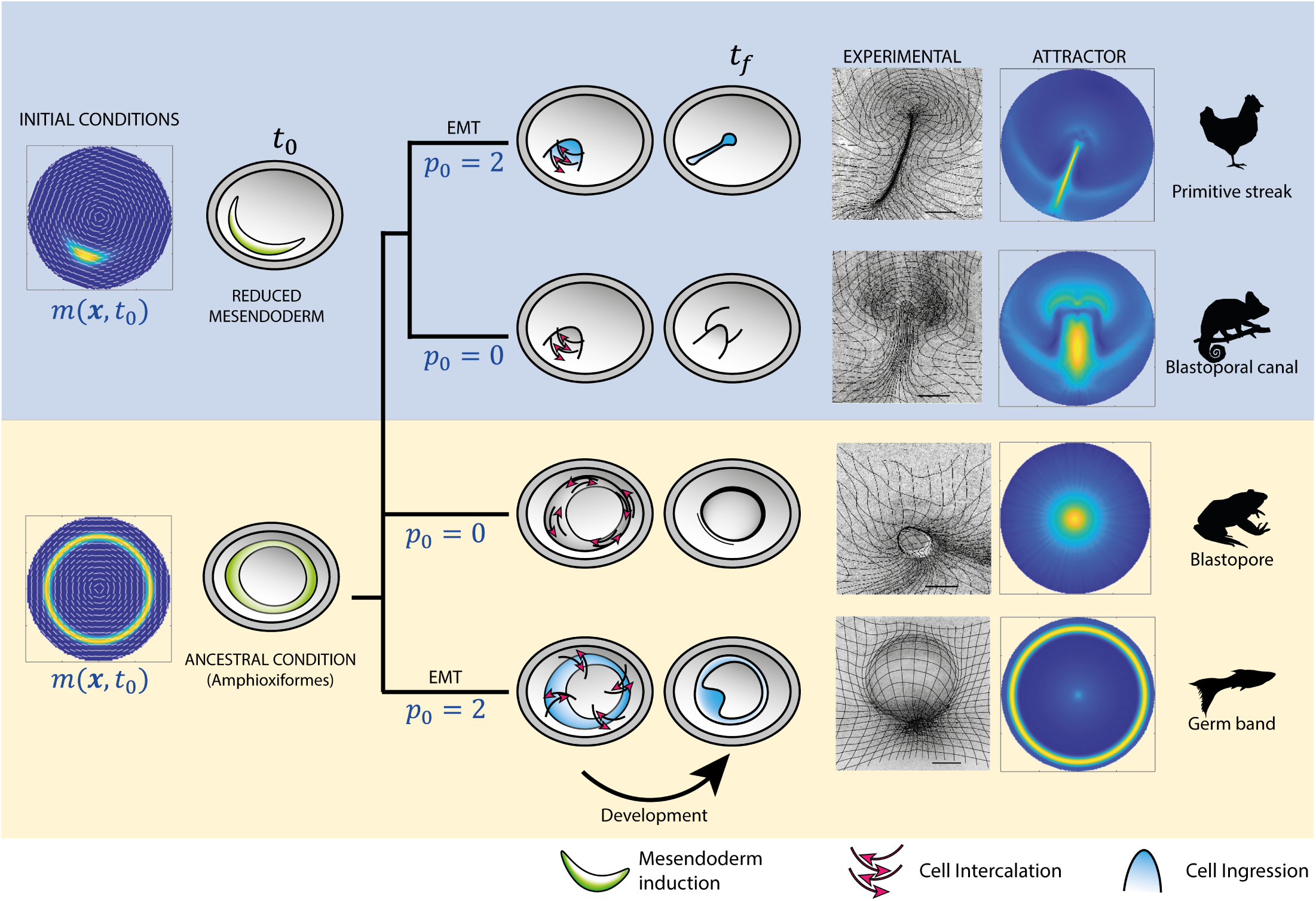
Evolutionary transitions in gastrulation patterns. Changes in a critical morphogenetic parameter *p*_0_ (the isotropic to anisotropic active stress ratio and regulates the amount of active cell ingression and EMT) and in the initial conditions *m*(*t*_0_) (the extent of the mesendoderm precursor territory) recapitulate major evolutionary transitions of vertebrate gastrulation, as reproduced experimentally in the chick embryo (see accompanying paper (9)). The right columns show deformed Lagrangian grids overlaid to the light-sheet microscope fields (9) and the attractors from Figs. 2-3 associated with different gastrulation modes. Scale bars are 500 *μm*.

## Acknowledgments

MS acknowledges the Schmidt Science Fellowship and the Postdoc Mobility Fellowship from the Swiss National Foundation for partial support. GSN acknowledges support from an EASTBIO BBSRC PhD student training grant (1785593). CJW thanks the BBSRC (BB/N009789/1, BB/K00204X/1, BB/R000441/1, BB/T006781/1) for financial support and a Wellcome Trust imaging equipment award (101468/Z/13/Z) for partial support. LM thanks the NSF-Simons Center for Mathematical and Statistical Analysis of Biology Award 1764269, NIH 1R01HD097068, the Simons Foundation, and the Henri Seydoux Fund for partial support.

## Supplementary Materials

### 1 Mathematical Model

We derive our model in Cartesian coordinates first, and then provide an equivalent reformulation in polar coordinates.

#### 1.1 Model in Cartesian coordinates

We denote the cell velocity field by **v**(**x**, *t*) = [*u*(**x**, *t*), *v*(**x**, *t*)]^*T*^, its vorticity scalar by *ω*(**x**, *t*), and we recall the velocity gradient decomposition

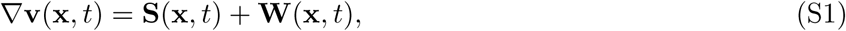

with the rate-of-strain tensor **S** and the spin tensor **W** defined as

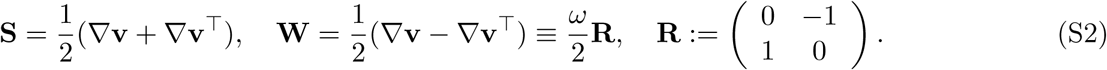

We further decompose **S** into its isotropic and deviatoric parts as

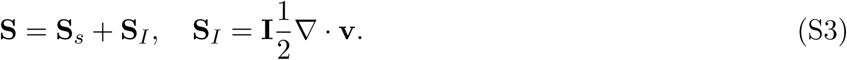

The viscous, active, total stresses and total stress trace are

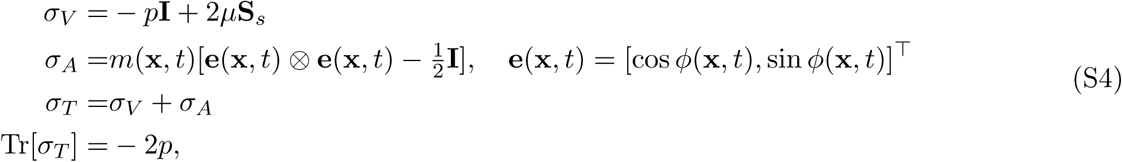

where *µ* denotes the shear viscosity, *m* denotes the stress magnitude arising from the active myosin concentration and *ϕ* the orientation of the myosin cables with respect to the *x*–axis. The active stresses tensor *σ*_*A*_ has eigenvalues ±*m/*2 and eigenvectors **{e, Re}**. In our model, a cell can be thought of as two springs in parallel: one representing the actomyosin cables that exert the active stress, and the second one representing the rest of the cell exerting the viscous stress. Therefore, the actomyosin cables tension can be computed as

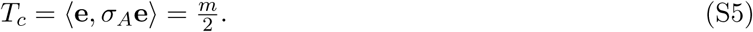

It is known that actin and myosin are characterized by catch bonds (1), i.e. that the dissociation rate of acto-myosin bonds depends on the tension along the actomyosin cable *T*_*c*_, and decreases exponentially for increasing tension. Using this assumption and eq. (S5), we model the evolution of active stress magnitude as

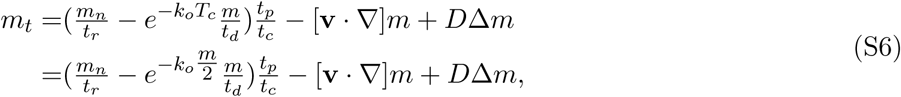

where *t*_*r*_ is the recruitment time scale of myosin from the cytoplasm, which has a given constant concentration *m*_*n*_, and 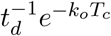 represents the tension-dependent dissociation rate of actomyosin cables. Note again that *m, m*_*n*_ have units [*Pa*] as they describe the stress magnitude arising from active myosin. From the time when active myosin is available to the time it generates active stresses, there is a time scale associated with the phosphorylation process (*t*_*p*_). We model this delay with the ratio (*t*_*p*_*/t*_*c*_), where *t*_*c*_ is the characteristic time scale of the system. Finally, the last two terms represent the advection term and the diffusion of active stress magnitude with diffusivity *D*.

We assume the evolution of actomyosin cables orientation satisfies Jeffery’s equation (2)

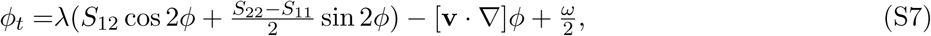

which describe the direction evolution of axis-symmetric fibers under the influence of a low Reynold’s number flow. In eq. (S7), 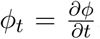, *λ* is a parameter quantifying the sensitivity of actomyosin cables reorientation due to shear-induced rotations, *S*_*ij*_ denotes the entries of **S** and the last two terms represent the advection term and the rigid-body rotation rate described by half of the flow vorticity.

Because viscosity is high (*µ* ≈ 9 × 10^3^ *Pa · s*), we ignore inertial terms and write the momentum balance as

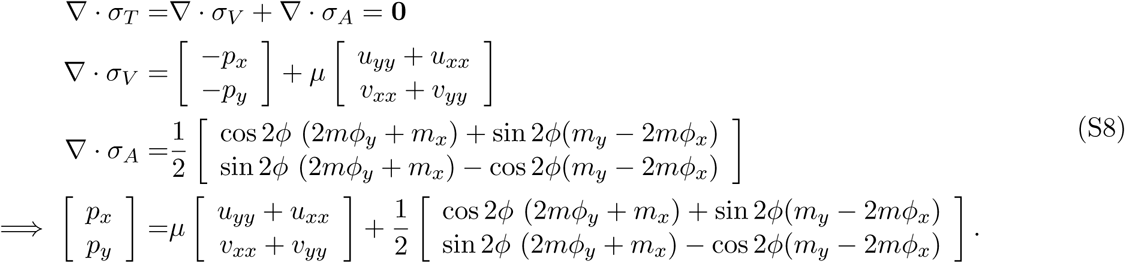

To close the system, we choose a simple continuity equation in which the flow divergence is proportional to the average total stress (eq. (S4)) and active myosin concentration as

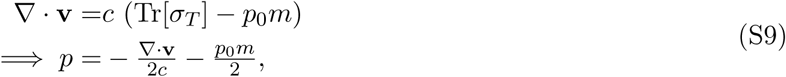

where *c* has units [*Pa* · *s*]^*−*1^ and *p*_0_ is a non-dimensional parameter modulating the contribution of active myosin to flow divergence and can be thought of as the ratio between the isotropic to anisotropic active stress. The intuition behind eq. (S9) is that regions of high compressive stress (averaged over all directions) and high active myosin exhibit negative flow divergence, i.e. cell ingression. Substituting eq. (S9) into eq. (S8), we obtain the resulting PDEs describing our active continuum without pressure terms

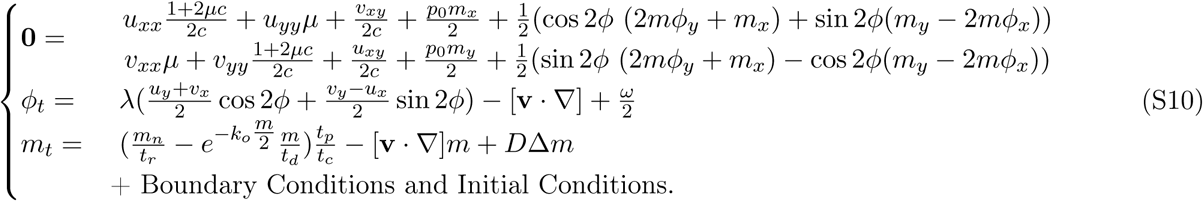

##### 1.1.1 Non-dimensional model in Cartesian coordinates

Denoting by *u*_*c*_, *x*_*c*_, *t*_*c*_ = *x*_*c*_*/u*_*c*_ and *m*_*c*_ = *µu*_*c*_*/x*_*c*_ the characteristic velocity, characteristic length scale, characteristic time scale and the characteristic viscous shear stress, we rewrite eq. (S10) in non-dimensional compact form as

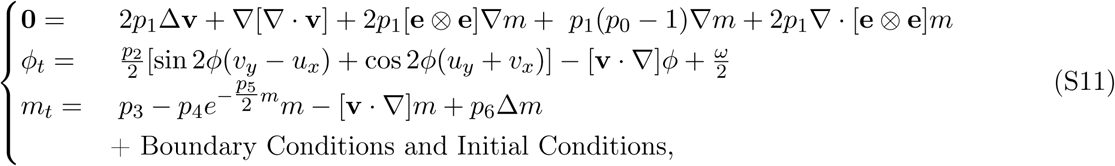

where *m, ϕ, u, v, x, y, t* are nondimensional. In eq. (S11), there are seven nondimensional parameters

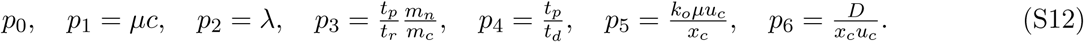

*p*_0_ describes the ratio of isotropic to anisotropic active stresses, *p*_1_ describes the ratio of the shear to bulk viscosity, *p*_2_ *∈* [0, 1] is the alignment parameter that describes how the actomyosin cables tend to orient due to shear-induced rotations, *p*_3_ represents the ratio of the myosin recruitment time scale (from the ambient myosin pool of concentration *m*_*n*_*/m*_*c*_) to the characteristic phosphorylation time scale, multiplied by *m*_*n*_*/m*_*c*_. *p*_4_ represents the ratio of the myosin phosphorylation time scale to the dissociation time scale. *p*_5_ is the product of the characteristic viscous shear stress and *k*_*o*_, which represents the inverse of the characteristic tension of the actomyosin cables (with units [*Pa*]^*−*1^). *p*_6_ is the inverse of the Peclet number characterizing the ratio between diffusive and advective transport of the active stress magnitude.

The first two terms of the force balance are a viscous force and a force arising from compressibility inhomogeneities. Instead, the last three terms are active forces induced by *m* and *ϕ* inhomogeneities. To gain insights on *p*_0_, we rewrite the force balance as

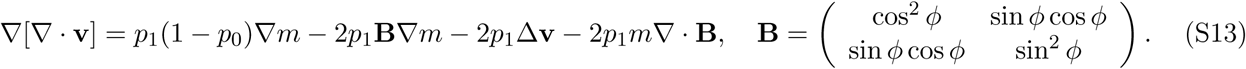

Assuming a uniform ridge of active myosin in the *y* direction, cables parallel to the ridge as in the case of perturbation 3 (Fig. 3E-F and Fig. 3 of (3), and neglecting the last two terms, we can see how *p*_0_ affects the relationship between the gradient of myosin and the one of the velocity divergence (SFig. 1). Specifically, when the isotropic to anisotropic active stress ratio *p*_0_ *>* 1, a ridge of high myosin induces a sink in the velocity field. By contrast, *p*_0_ *<* 1 induces a source. Comparing our reasoning with experimental observations (Fig. 1L and Fig. 3 of (3)), in our model we use *p*_0_ *>* 1 or *p*_0_ = 0.

**Supplementary Figure 1:**
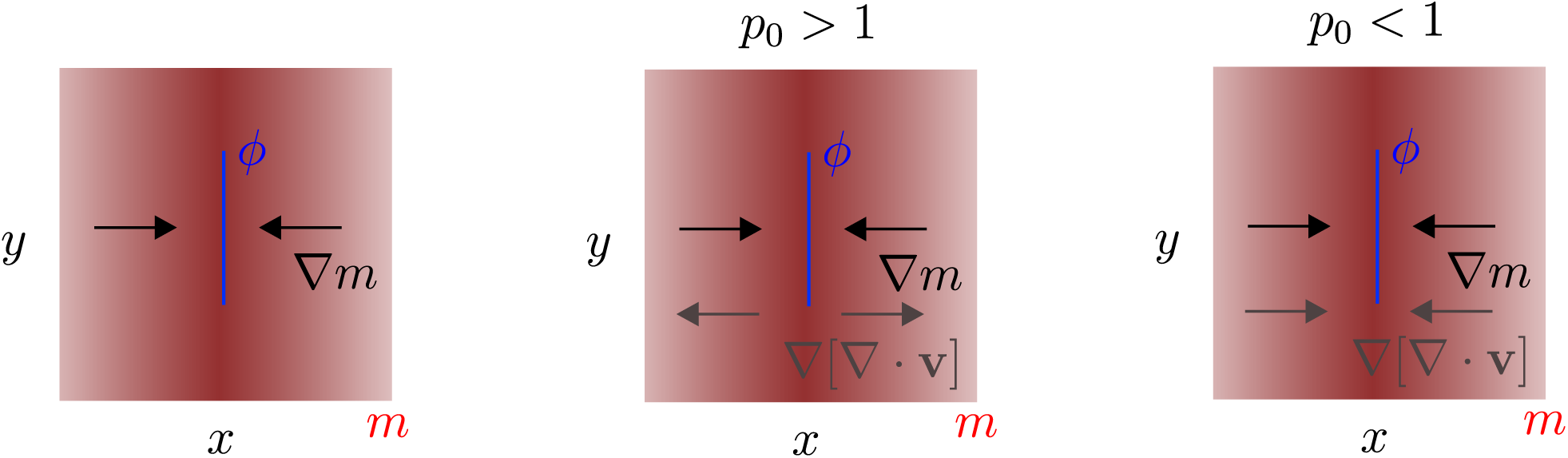
Influence of *p*_0_ on the relationship between ∇*m* and ∇ [∇ ·v]. We consider only the first three terms of eq. (S13) and a simple distribution *m* made of a ridge in the *y* direction with actomyosin cables parallel to the ridge.

Finally, observing the first two terms in the last equation in (S11), we note that more myosin induces higher tension on the cables, which in turn decreases the dissociation rate: (*m* ↑) → *Tc* ↑ (more positive) → diss. rate ↓ → myosin increases (*m* ↑) (Fig. 2A). This observation suggests an instability, which will become clear in the next section.

##### 1.1.2 One-dimensional model in Cartesian coordinates

To gain insights about model (S11), we assume slow dynamics in the *y−* direction, ignore the *ϕ−* dynamics and set *ϕ* = 0, obtaining the following simplified one-dimensional model

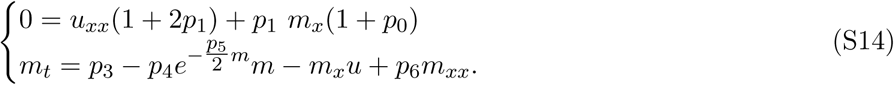

Because in 1D there is no difference between isotropic and anisotropic stress, we set *p*_0_ = 0 without altering the relevant properties of our simplified model. Equation (S14) describes the dynamics along a straight line perpendicular to the primitive streak. We set Dirichlet boundary conditions on the velocity *u*(0, *t*) =*−*|*u*_*b*_|, *u*(*L, t*) = |*u*_*b*_| consistent with the symmetric cell migration in the extra-embryonic area, and no-flux boundary conditions for *m*: *m*_*x*_(0, *t*) = *m*_*x*_(*L, t*) = 0.

We investigate the main features of eq. (S14) by seeking the constant fixed points for *m* which solve the implicit equation 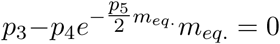. From the first equation, we compute the corresponding equilibrium velocity field 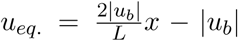. We then study the linear stability of these equilibrium solutions by computing the spectrum of the corresponding linearized PDE. Denoting by *δ***h** = [*δu, δm*]^*T*^ the perturbation from the equilibrium, we compute the linearized version of Eq. (S14) in operator form as 𝒜*δ***h** = ℬ (*x*)*δ***h** where

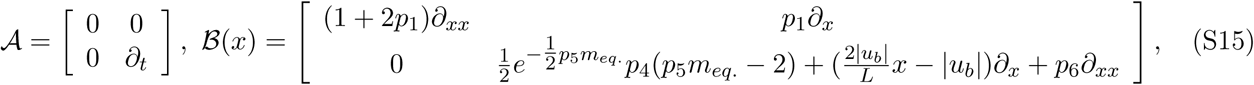

whose boundary conditions are inherited from the nonlinear problem, and correspond to zero Dirichlet boundary conditions for *δu* and zero flux boundary conditions for *δm*. We discretize the linearized PDE using a centered finite differencing scheme (4) and compute the generalized eigenvalue problem associated with eq. (S15).

**Supplementary Figure 2:**
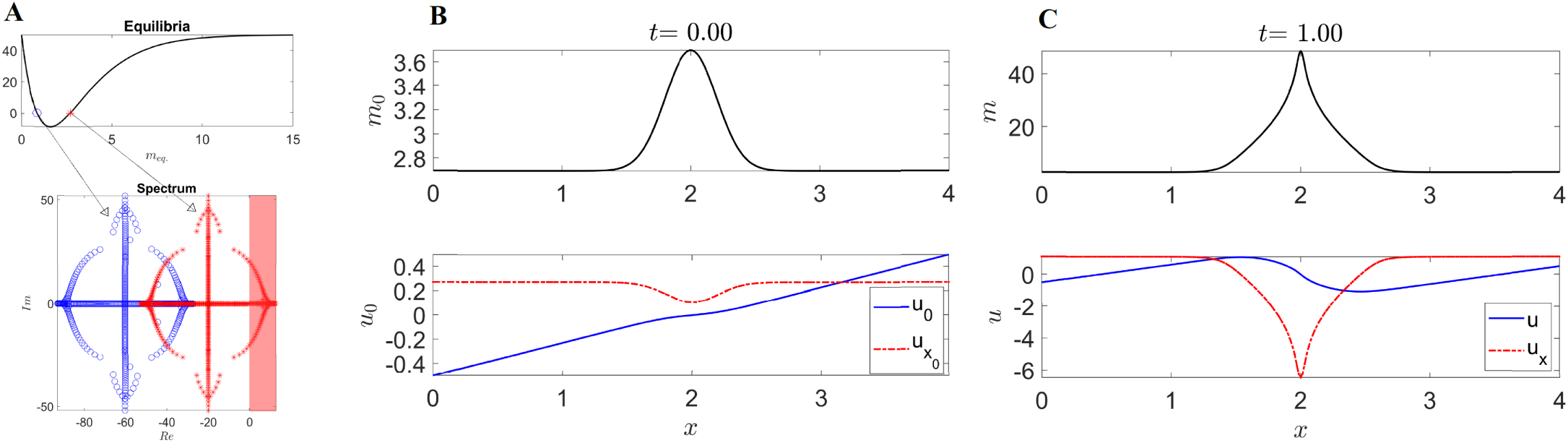
1D model. (A) Top: Graph of 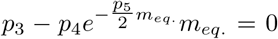 shows two *m*_*eq*._ marked with a blue circle and a red asterisk. Bottom: Truncated spectrum of the linearized PDE around *m*_*eq*._, *u*_*eq*._, consisting of the 500 eigenvalues with the highest real part. (B) Initial condition of the nonlinear 1D model using as initial myosin concentration a Gaussian added to the higher unstable *m*_*eq*._. The blue curve shows the associated initial velocity *u*_0_, and the red dashed curve the initial velocity divergence 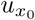. (C) Same as B at the final time. The full time evolution is available as [Movie 1](https://www.dropbox.com/s/6pgxys02xe2or9e/1D%20Model.mp4?dl=0). Parameters: *p*_0_ = 0, *p*_1_ = 0.25, *p*_3_ = 100, *p*_4_ = 100, *p*_5_ = 1.25, *p*_6_ = 0.001, *u*_*b*_ = 0.5, *L* = 4, *dx* = 0.008, *dt* = 0.0001.

Supplementary Figure 2A top shows two constant equilibria *m*_*eq*._, while the bottom panel shows the corresponding 500 eigenvalues with the largest real part of the linearized operator for both the equilibria. The spectrum suggests that the low (high) myosin concentration is linearly stable (unstable) for the chosen parameter values. We confirm these linearized results by solving the full nonlinear PDE from an initial Gaussian myosin distribution added to the unstable *m*_*eq*._ (SFig. 2B top). From a biological perspective, a region of higher *m*_0_ represents the mesendoderm precursor. At later times (SFig. 2C), *m* undergoes a focusing instability, that in turn generates a 1D attractor corresponding to the PS and marked by a peak of negative divergence (red dashed curve). The complete time evolution is available as [Movie 1](https://www.dropbox.com/s/6pgxys02xe2or9e/1D%20Model.mp4?dl=0). Finally, the velocity divergence *u*_*x*_ shows a contracting region close to the PS as well as two symmetric expanding regions close to the boundary, consistent with the isotropic deformations observed in the Embryonic and Extra-Embryonic territories (5). Our simple 1D model reproduces several key features of amniote gastrulation and perhaps elucidates some of its hidden driving mechanisms. Yet, it is insufficient to reproduce the convergent extension and vortical patterns observed in experiments, which we investigate with our 2D model.

#### 1.2 Model in polar coordinates

Because of the circular geometry of the Extra-embryonic tissue as well as the symmetric radial cellular motion at its outer boundary, we re-derive our model (S10) in polar coordinates.

**Supplementary Figure 3:**
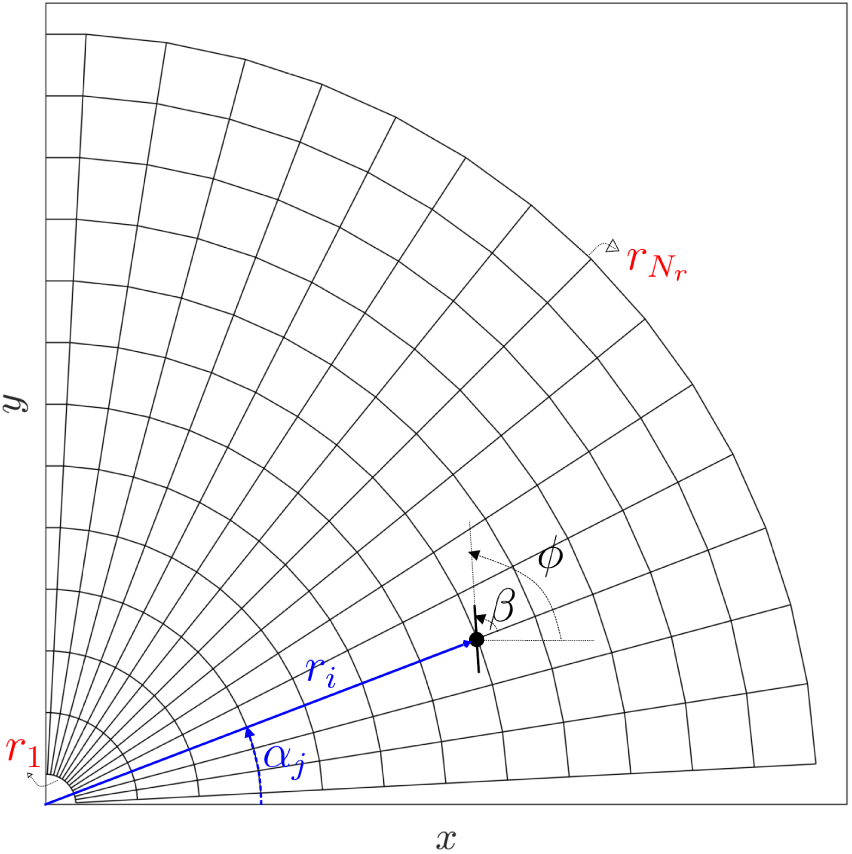
Numerical mesh. Illustration of the discretization grid (first quadrant) used to solve eq. (S17). *β* denotes the cables orientation relative to *α*.

Denoting the velocity field by **v** = *u*(*r, α, t*)**e**_**r**_ + *v*(*r, α, t*)**e**_*α*_ and the cables orientation relative to *α* by *β* (SFig. 3), we obtain

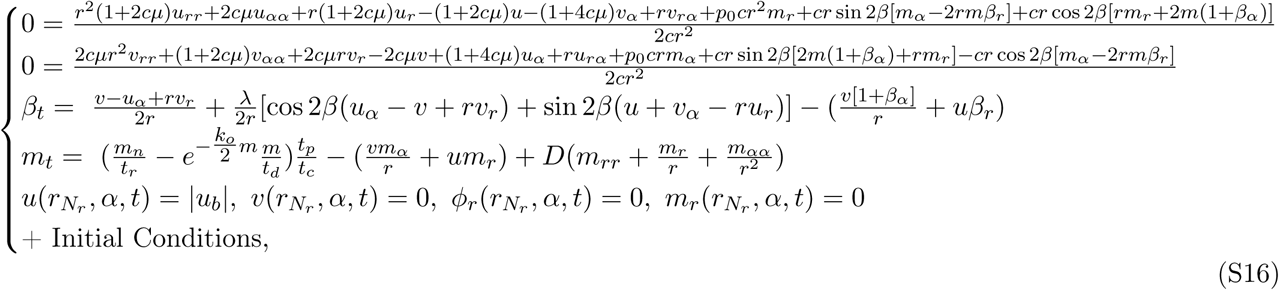

which, in non-dimensional form, becomes

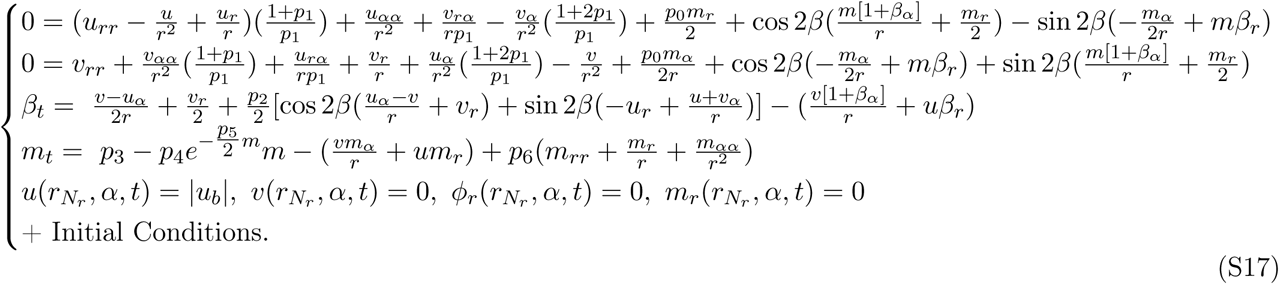

### 2 Parameters

In our simulations, we select the parameter values as follows. *p*_0_ = 0 when active cell ingression is inhibited with drugs and *p*_0_ = 2 otherwise. We set *p*_1_ = 0.15 and note that increasing *p*_1_ models a more compressible tissue. With higher *p*_1_, the typical counter-rotating vortices (polonaise movement) disappear due to increased cell ingression and less cell redistribution. We set *p*_2_ = 0.9 as we model actomyosin cables as elongated passive fibers. To get the order of magnitude of *p*_3_, *p*_4_, we note that the phosphorylation time scale (*t*_*p*_) to generate active stress from active myosin is ≈ 20min, while the recruitment and dissociation time scales *t*_*r*_ ≃ *t*_*d*_ ≈ 10s. With these estimates, we set *p*_3_ = 50 and *p*_4_ = 100. We are unaware of techniques to estimate *p*_5_, and set *p*_5_ = 1.25. We find, however, that slight variations in *p*_3_, *p*_4_, *p*_5_ do not alter the overall system behavior as long as the topology of the graph (SFig. 2A) determining *m*_*eq*._ does not change significantly. We set *p*_6_ = 0.001 and note that its variations have negligible effects on our results. Finally, because *∂*Ω (Fig. 1A) is close to the EP-EE boundary, where the normal velocity is zero, we set |**v**_b_| = 0.1. Slight changes in **v**_b_ do not affect the qualitative behavior of our results. We summarize our parameters in table 1.

**Table 1:** Parameter values.

| $p_0$ | $p_1$ | $p_2$ | $p_3$ | $p_4$ | $p_5$ | $p_6$ |
| --- | --- | --- | --- | --- | --- | --- |
| 0 or 2 | 0.15 | 0.9 | 50 | 100 | 1.25 | 0.001 |

Overall, we have selected parameter values using a combination of mechanistic arguments and numerical tests. A detailed analysis of the parameter space is planned for future work.

### 3 Numerical Scheme

We develop a finite-difference numerical scheme to solve eq. (S17) in MATLAB. First, we multiply the momentum balance equations by *r*^2^, leaving their solutions unaltered while avoiding numerical issues at grid points close to the origin. Then, we discretize spatial differential operators using centered finite-difference at second-order accuracy, except at the grid points (*r*_1_, *α*_*j*_) (SFig. 3), where we use forward finite-difference at second-order accuracy (4). We solve the first two equations for *u, v* by inverting (once) the corresponding discretized differential operator, which we multiply at each time step to the discretized forcing vector arising from the updated *β, m*-dependent terms. We adopt a two-step Adams–Bashforth method to solve the last two PDEs for *β* and *α*.

Because the cables orientation is a direction instead of a vector field, spatial derivatives of *β* need special care. Specifically, when computing 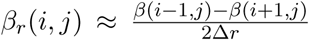, one should make sure that both the sign and the amplitude are correct. We do so by rewriting 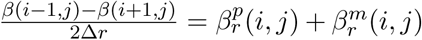, where 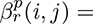 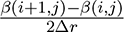, 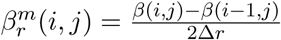, and compute 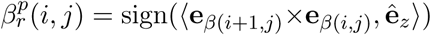 arccos, ⟨**e**_*β(i+1,j)*_, **e**_*β(i,j)*_⟩ where **e**_*β*_ = [cos *β*, sin *β*, 0], 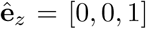. The same strategy can be used for higher-order derivatives and forward finite difference schemes. We summarize our numerical scheme in the following algorithm.

#### Algorithm 1 Numerical Solver of eq. (S17)

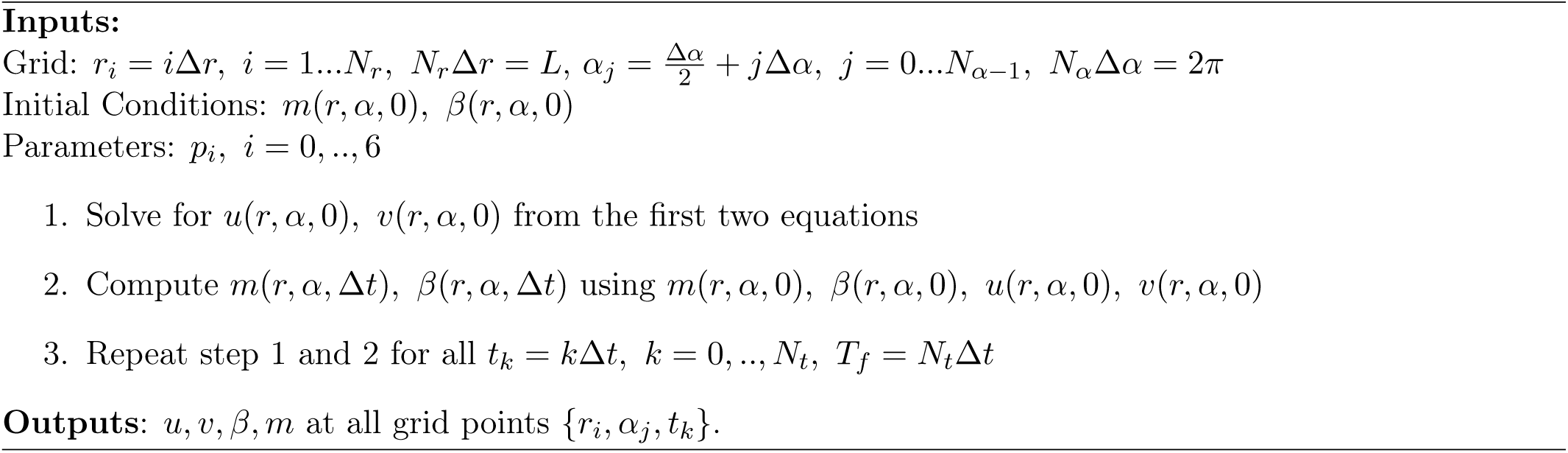

To prevent the development of sharp derivatives in *β* at the origin due to the forward finite-difference scheme, we add a diffusion term in the *β*_*t*_ equation with a non-dimensional prefactor of 10^−3^. In all our simulations, we use Δ*r* = 0.05, *L* = 1, Δ*α* = 4°, Δ*t* = 5 × 10^−6^, *T*_*f*_ = 1. We monitor the accuracy of our numerical solver by substituting our solution at each time step into the momentum balance equations. We find deviations from zero of the order ≈ 10^−11^.

### 4 Dynamic Morphhoskeletons

Given a modelled or experimental planar velocity field **v**(**x**, *t*), we compute the Dynamic Morphoskeleton (DM) (6) from the backward and forward Finite Time Lyapunov Exponents (FTLE). We compute the FTLE

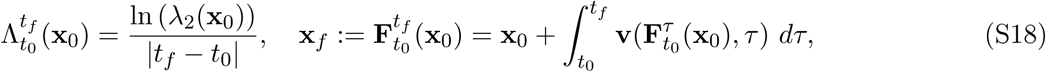

where *λ*_2_(**x**_0_) denotes the highest singular value of the Jacobian of the flow map 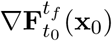 and 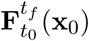 the flow map describing the trajectories from their initial **x**_0_ to final **x**_*f*_ positions. To compute the FTLE, we first calculate 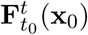 by integrating the cell velocity field **v**(**x**, *t*) using the MATLAB built-in Runge-Kutta solver ODE45 with absolute and relative tolerance of 10^−6^, linear interpolation in space and time, and a uniform dense grid of initial conditions. Then, denoting the *i* − *th* component of the flow map 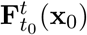 by 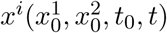, we compute the deformation gradient 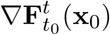 using the finite-difference approximation

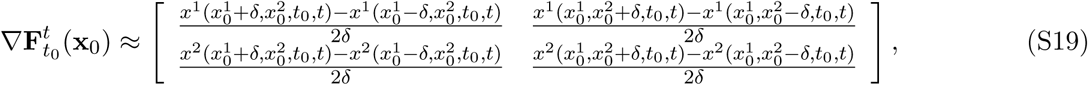

where *δ* is the initial conditions’ grid spacing. After computing 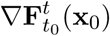, we use eq. (S18) for computing the FTLE field.

## 5 Supplementary figures

**Supplementary Figure 4:**
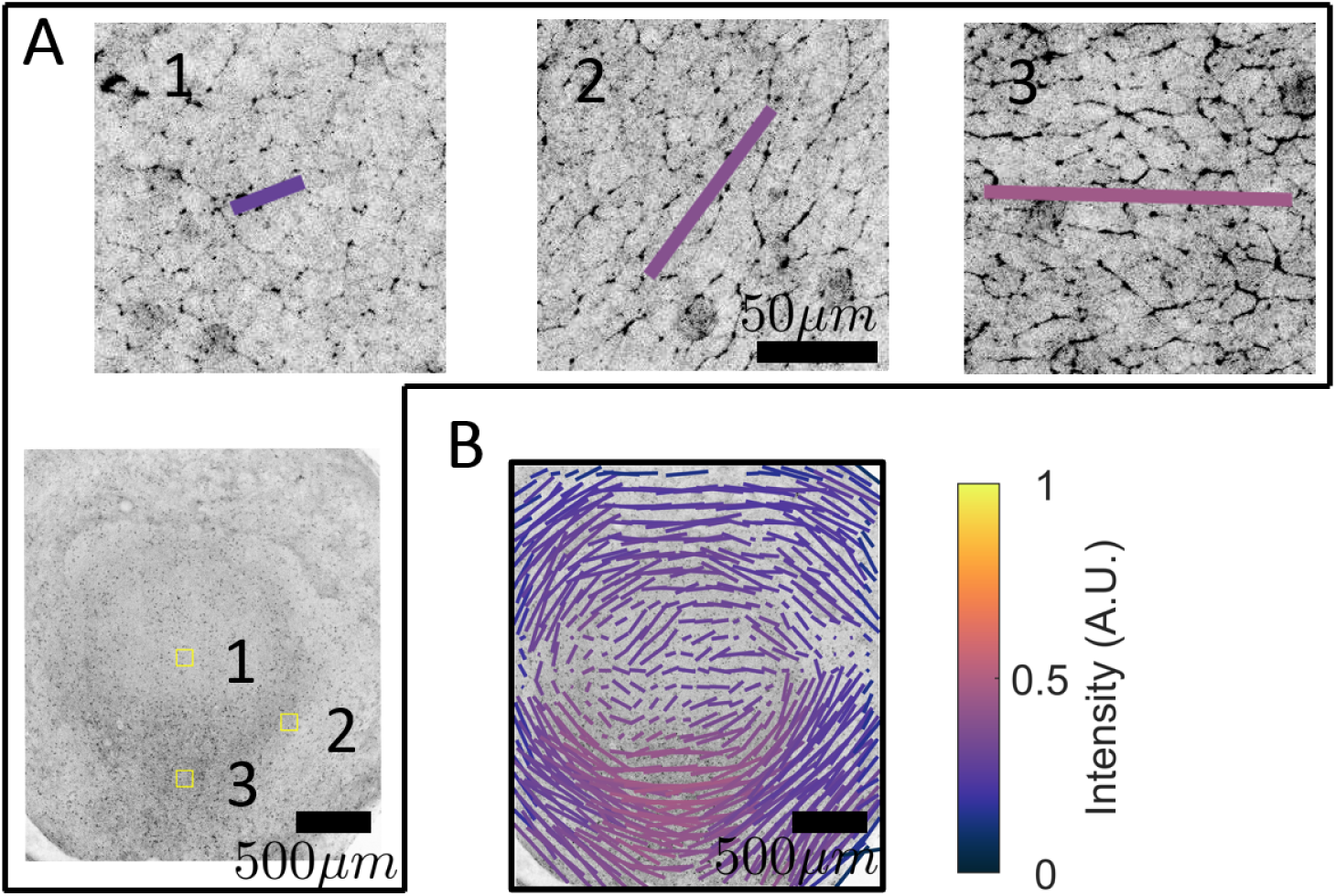
Active myosin cables. A) Phospho-myosin light chain (pMLC) expression in the chick gastrula (HH3). The panels 1-3 are taken at the positions of the yellow squares in the overview image show the formation of pMLC cables spanning several cells in panels 1 and 2. The magnitude and direction of the bars in panels 1-3 quantify the anisotropy of the pMLC cables, the color of the bars indicate the relative pMLC concentration. B) The pattern of pMLC cables of the HH3 stage embryo shown in A. See the accompanying paper (3) for details on the quantification of pMLC cables.

**Supplementary Figure 5:**
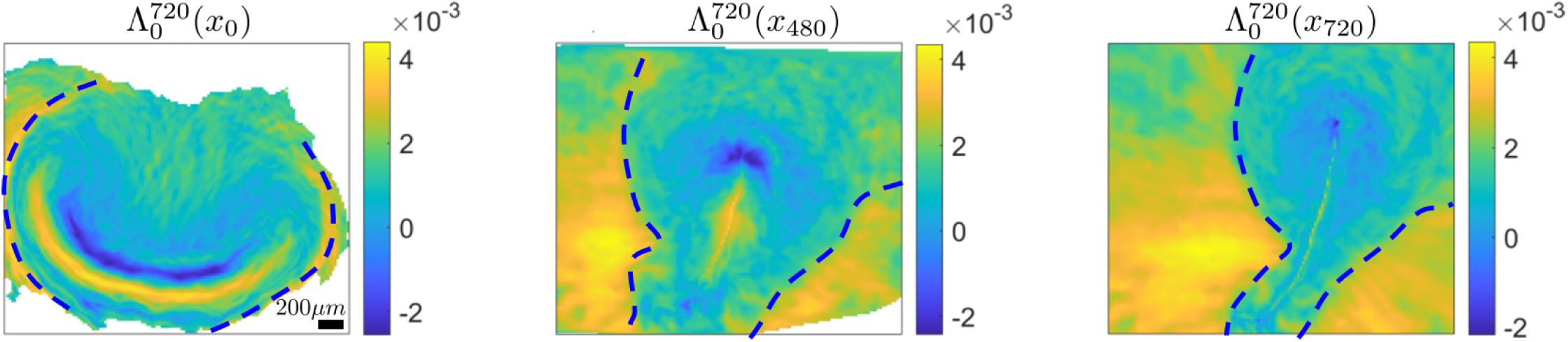
Dynamic EE-EP repeller in wild-type chick gastrulation. Left) High values of 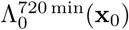 mark two repellers displayed at **x**_0_. The dashed curve marks the EE-EP repeller. Center) Position of the EE-EP repeller after 480min along with the 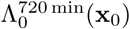 visualized on the current cell position 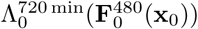. Right) same as left but after 720min. [SM Movie1](https://www.dropbox.com/s/yslitzz0wwjon92/OuterRepEvolution_CTR0245_M2Exp.mp4?dl=0) shows the corresponding time evolution. Colorbars show attraction rates in min^*−*^.

